# VirEvol platform : accurate prediction and visualization of SARS-CoV-2 evolutionary trajectory based on protein language model, structural information and immunological recognition mechanism

**DOI:** 10.1101/2023.09.15.557978

**Authors:** Xincheng Zeng, Linghao Zhang, Zhenyu Ning, Yusong Qiu, Ruobing Dong, Xiangyi Li, Lijun Lv, Hanlin Xu, Yanjing Wang, Buyong Ma

## Abstract

Predicting the mutation direction of SARS-CoV-2 using exploratory computational methods presents a challenging, yet prospective, research avenue. However, existing research methods often ignore the effects of protein structure and multi-source viral information on mutation prediction, making it difficult to accurately predict the evolutionary trend of the SARS-CoV-2 S protein receptor-binding domain (RBD). To overcome this limitation, we proposed an interpretable language model combining structural, sequence and immune information. The dual utility of this model lies in its ability to predict SARS-CoV-2’s affinity for the ACE2 receptor, and to assess its potential for immune evasion. Additionally, it explores the mutation trend of SARS-CoV-2 via a genetic algorithm-directed evolution. The model exhibits high accuracy in both regards and has displayed promising early warning capabilities, effectively identifying 13 out of 14 high-risk strains, marking a success rate of 93%.”. This study provides a novel method for discerning the molecular evolutionary pattern, as well as predicting the evolutionary trend of SARS-CoV-2 which is of great significance for vaccine design and drug development of new coronaviruses. We further developed VirEvol, a unique platform designed to visualize the evolutionary trajectories of novel SARS-CoV-2 strains, thereby facilitating real-time predictive analysis for researchers. The methodologies adopted in this work may inspire new strategies and offer technical support for addressing challenges posed by other highly mutable viruses.

## Introduction

### Research Background

Currently, the SARS-CoV-2 pandemics is still evolving and spreading globally, and the proliferation of virus variants poses a great threat to human life and health and global economic development. While most mutations reduce the overall fitness of the virus, some mutations, individually or in combination, result in high-risk variants (HRVs) with altered immune evasion or increased transmissibility [1]. The mutation sites of SARS-CoV-2 mutants are mainly focused on structural proteins, especially on the spike (S) protein [2]. The S protein is responsible for the binding and membrane fusion of the virus to the host cell membrane receptor of angiotensin-converting enzyme 2 (ACE2), and it is an important site for the neutralizing antibody of the host as well as a key target for vaccine design [3].

### Current Research progress in China and abroad

Early studies have focused on discovering mutational roles and effects through high-throughput sequencing of RBDs [4, 5, 6]. Starr et al. [6] and Greaney et al. [4] performed deep mutation scanning (DMS) experiments on the SARS-CoV-2 S-protein RBD sequences to determine the effects of substitutions at a single site on the binding ability with ACE2 receptor and monoclonal antibodies. However, wet experiments are time and resource consuming and cannot monitor the vast protein variant sequence space.

Some researchers have started using machine learning methods to construct computational models to obtain possible effects of mutations on protein function. Alexander et al. [7] trained a large-scale Transformer model with a self-supervised protein language modeling goal to infer the effects of mutations without supervision. Chloe et al. [8] combined a linear regression with a Potts model to obtain a data-efficient model for adaptive inference of variants. These models rely only on sequence data and have a simpler model structure, which often lead to poor prediction accuracy.

In terms of viral affinity prediction, Chen et al. [9] focused on using molecular dynamics (MD) simulations combined with neural networks to predict RBD binding with ACE2 receptor, with a correlation of 0.73 when compared to binding affinity measurements from DMS. Wang et al. [10] used a neural network to directly learn RBD-ACE2 receptor binding from DMS data without the need for MD simulations, obtained a significant improvement of model accuracy. Abbasi et al. [11] worked on developing a model for predicting protein-protein binding based only on sequence, and their team proposed a novel sequence-based protein binding affinity predictor, ISLAND. However, the higher computational cost of MD simulation for structural optimization has limited its wide application in current research.

To predict the immune escape ability of the virus, Brian Hie [12] proposed a neural network model that was trained to predict the escape rate of SARS-CoV-2 RBD when bound to any antibody using deep mutation scanning experimental data. Eric Wang [13] further trained on 315,000 data points from deep mutation scanning experiments in order to improve the correlation coefficients, which allowed the model to achieve Spearman’s correlation coefficients of 0.46 and 0.52 on two reserved test sets. In addition, Yisimayi et al. [14] used DMS to characterize antigenic epitopes and showed that these Omicron-specific antibodies target at different RBD epitopes compared to those induced by wild type, as well as identified evolutionary hotspots of the XBB.1.5 RBD and predicted the direction in which the virus might mutate.

In recent years, neural network language models (NNLMs) based on recursion and attention mechanism have been used to learn biological language and design, which can be used not only for protein variant analysis [7, 15], but also to capture the effects of mutations on protein function [16]. Inspired by these works, Hie et al. [12] were able to predict the potential risk of COVID variants with multiple mutations by training a protein language model on a set of evolutionarily relevant sequences; Karim et al. [17] further combined protein language model scores with structural modeling to monitor the risk of existing variants. However, these approaches rely on existing data and they do not predict detailed pathways for the evolutionary potential of variants and antigens. Taft et al. [18] further performed deep learning of RBM sequences and developed a predictive model that was able to predict class I, II, and III antibodies for the SARS-CoV-2 variant in terms of ACE2 receptor binding and antibody escape, however, the model only focuses on the RBD region of a small portion of the RBD region and does not consider class IV antibodies.

The use of large-scale language models to predict the direction of mutation in SARS-CoV-2 is a promising research direction; however, current studies have several major limitations: first, the predictions of the language models are limited by the quality and diversity of the input data. The limited availability of actual mutation data for SARS-CoV-2 leads to bias or inaccuracy of the models in their predictions. Second, language model predictions are often based on statistics and pattern recognition and lack detailed explanations of underlying mechanisms. Finally, language models may ignore certain important factors, such as protein structure, immunological information, viral replication kinetics, and other key factors when making predictions, so relying on language model predictions alone may not provide a comprehensive understanding of the mutation trends of SARS-CoV-2.

In summary, an interpretable language model that can take account of protein structure and multiple sources of viral information is needed to accurately predict the evolutionary trend of the RBD region of the SARS-CoV-2 S protein.

### Project Program

To address the huge challenge of evolutionary prediction of a highly mutable virus (SARS-CoV-2), we collected more than 1.6 million sequence data, close to 40,000 deep mutation scans, and 594 high-quality virus-antibody complex structural data, and used a protein language model with 650 million covariates for sequence feature extraction and downstream task training to predict the affinity of SARS-CoV-2 with the target ACE2 receptor. VirEvol is an evolutionary prediction platform we built for SARS-CoV-2, which can reveal the molecular evolutionary patterns and predict the evolutionary trends of SARS-CoV-2, and help for vaccine design and drug applications against important SARS-CoV-2 mutation sites and potentially high-risk strains.

### Research method and prediction algorithm

#### Data collection

We collected nearly 1.65 million sequences of the novel coronavirus S protein, over 50,000 deep mutation scanning data for the novel coronavirus S protein, as well as high-quality data comprising 873 novel coronavirus antibodies and 1,478 novel coronavirus S protein structures. These data underwent thorough cleaning processes.

For sequence data, we removed duplicate sequences, sequences with amino acids labeled as “X,” and sequences with a length less than 1000. Regarding deep mutation scanning data, we excluded data with missing affinity values and duplicates. For structural data, we selected matching structures from the novel coronavirus antibody and novel coronavirus S protein structure datasets for structural recognition. Further details of data processing can be found in the supplementary materials.

#### Affinity prediction model

The affinity prediction model employs the most efficient protein language model architecture based on Transformer, as illustrated in Figure 2. We retrained the ESM-2 model with 650 million parameters and the ProBERT-BFD model with 420 million parameters.

**Figure 2.**
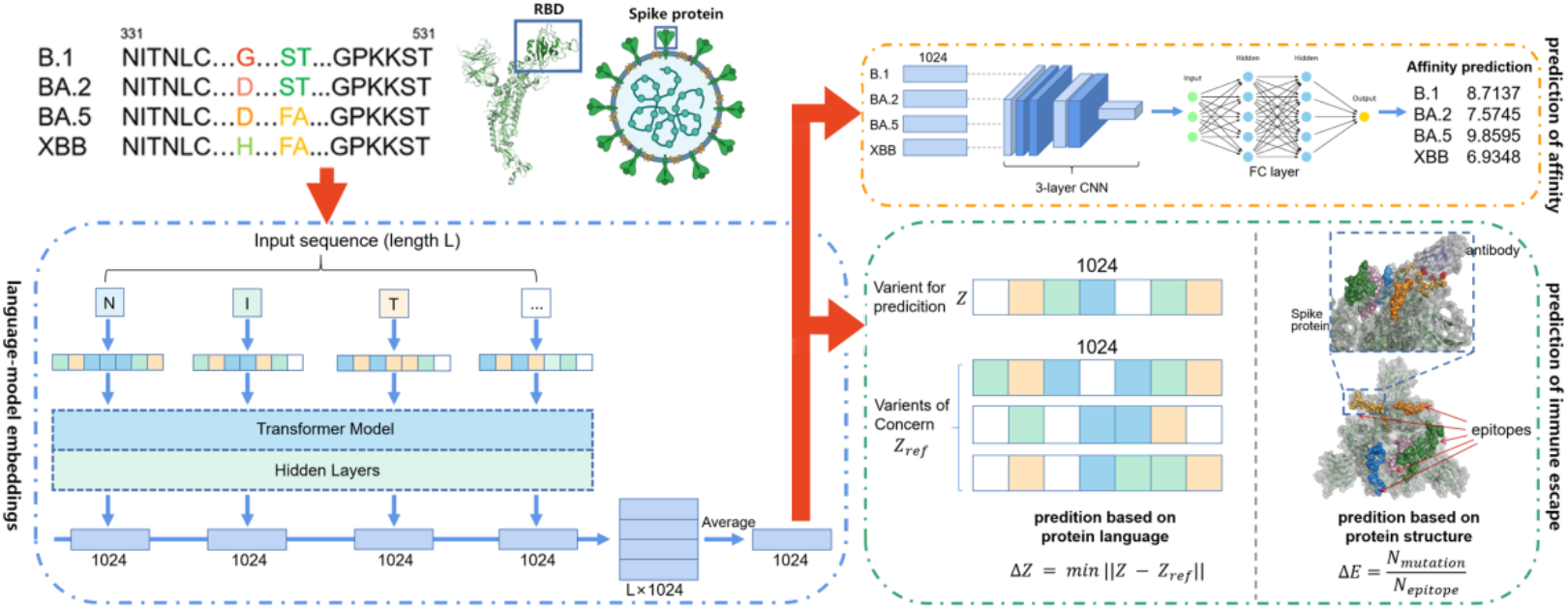
VirEvol Model Framework as a Predictive Platform for Molecular Evolution of the Novel Coronavirus

**Figure 3.**
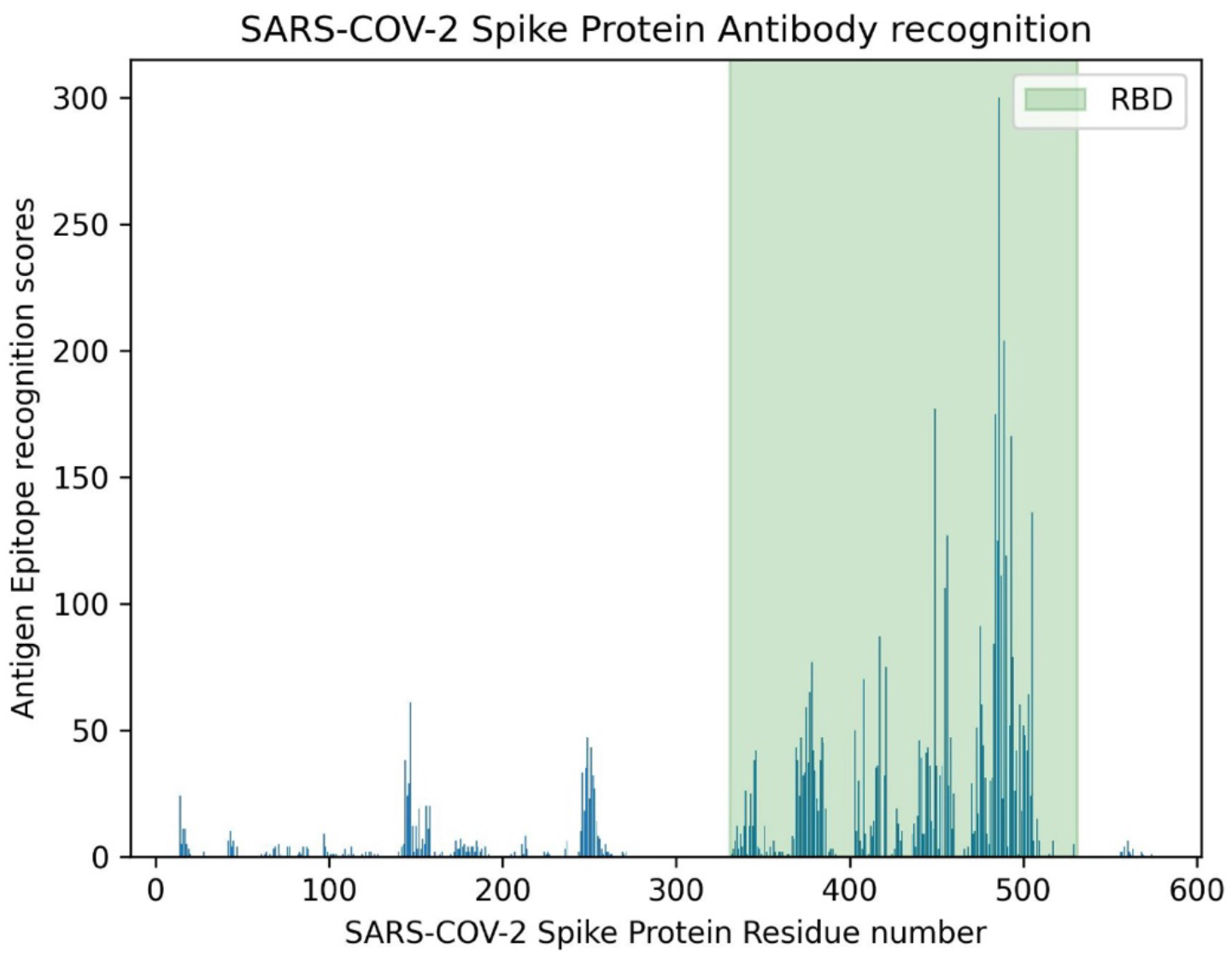
Important residue hotspots for COVID-19 antibody recognition

#### Immune Evasion Prediction Model

The immune evasion prediction score consists of structure-based immune evasion scores and language model-based immune evasion scores.

To calculate the structure-based immune evasion scores, we conducted three-dimensional structural scans on 594 antibody-antigen complexes, which covered antibodies from classes one through four, and were processed after downloading from CoV3D database[19]. From a structural perspective, we computed residue contacts between all antibodies and antigens and tallied important residue hotspots recognized by novel coronavirus antibodies to calculate the immune evasion scores using previously developed program[20].

To calculate the language model-based immune evasion scores, we utilized semantic vectors (embeddings) extracted from a protein language model to measure the degree of semantic change. For viruses, the reason for immune evasion lies in changes occurring in the virus’s antigenic epitopes. While maintaining the structural-functional foundation necessary for the virus to invade its target, there is an overall alteration in the virus’s semantic expression, allowing it to evade antibody recognition sites. Consequently, changes in semantic vectors can be used to compute immune evasion scores *ΔZ*.

#### Genetic evolution model

In the Genetic Evolution module, we perform virus adaptability calculations and multiple rounds of genetic variation iterations to screen potential high-risk strains, as shown in Figure 4. Unlike multi-round scans based on single-site saturation mutations, this method exponentially reduces the sequence search space, making it more efficient in exploring the boundaries of virus evolution. The virus adaptability prediction score is based on target affinity and immune evasion score scores, calculated according to the following formula:

**Figure 4.**
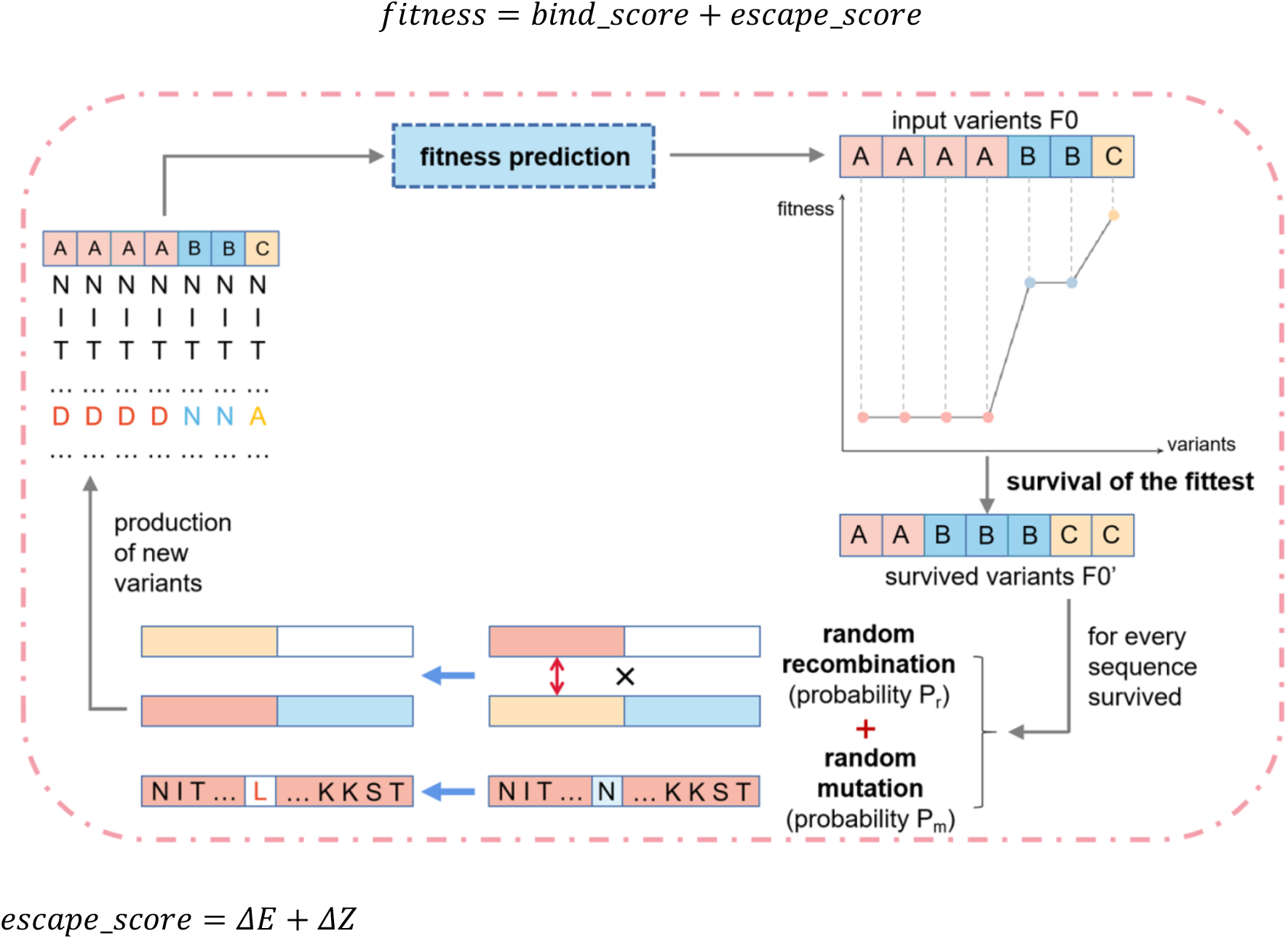
VirEvol as a Simulated System for the Evolution of the Novel Coronavirus

## Results

### Affinity prediction model achieves high accuracy

After fine-tuning, the accuracy of the protein language models with two different architectures was significantly improved in the task of predicting the affinity of the novel coronavirus ACE2 receptor, among which the predicted and true values *R*^2^ of ESM-VirEvol exceeded 0.9 on the training set and reached 0.88 on the validation set. The predicted and true values of the ProtBERT-VirEvol model approach 0.95 on the training set and exceed 0.85 on the validation set. Since the ESM-VirEvol model is more accurate on the validation set, it is used as the actual call model for VirEvol.

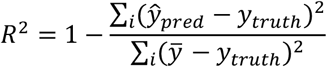

### Adaptive prediction of viral evolutionary direction based on affinity and immune escape ability

The model calculated adaptive indicators, which consist of viral affinity and immune escape ability, and made predictions on 39 mutation subtypes of COVID-19 from January 2020 to May 2023. The Spearman correlation between the earliest detection time of different COVID-19 mutants and the predicted adaptability index was more than 0.9, which accurately replicated the evolutionary trajectory of the virus, as shown in Figure 9. This indicates that the fitness algorithm has learned the key mechanism of virus evolution and can effectively identify the evolutionary trend of different strains.

**Figure 5.**
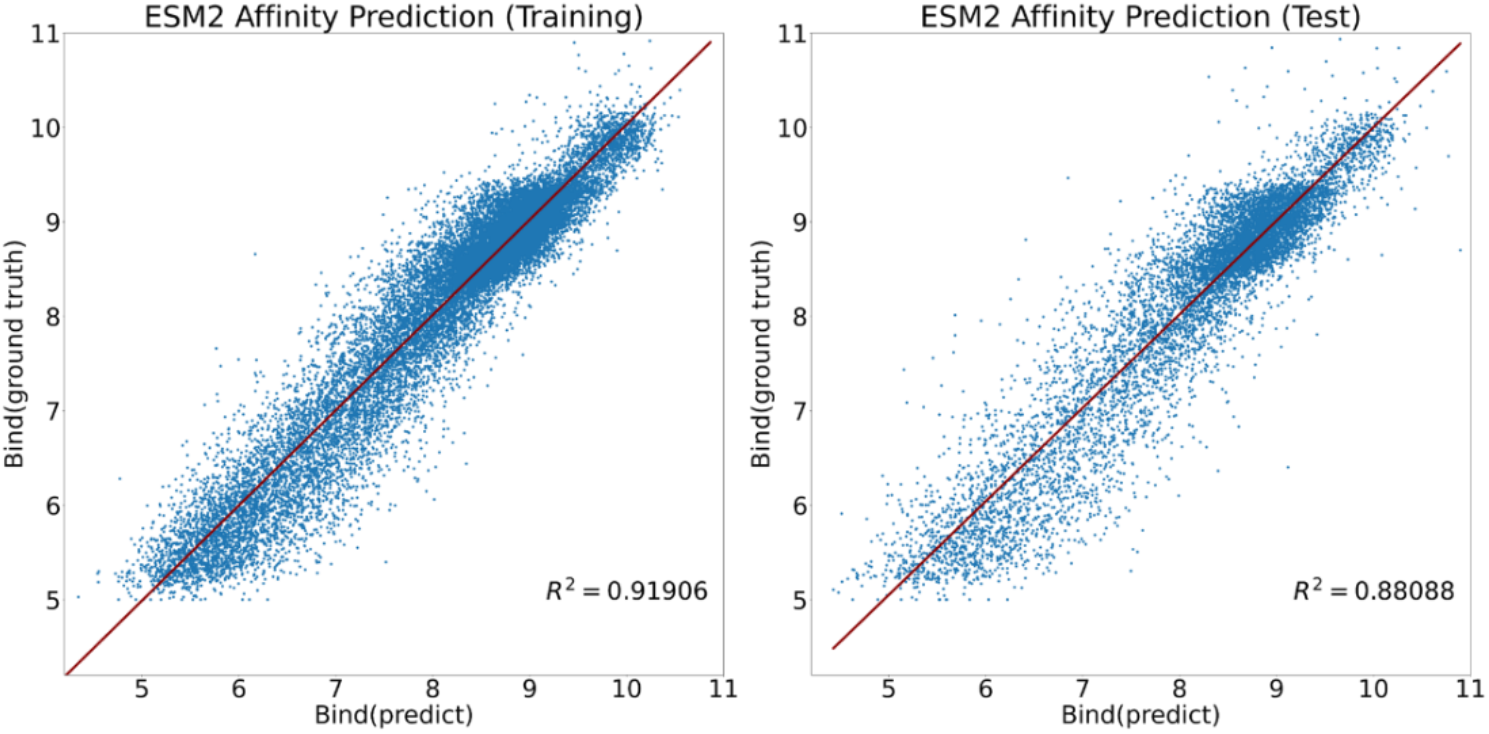
Affinity prediction results of the ESM-VirEvol model

**Figure 6.**
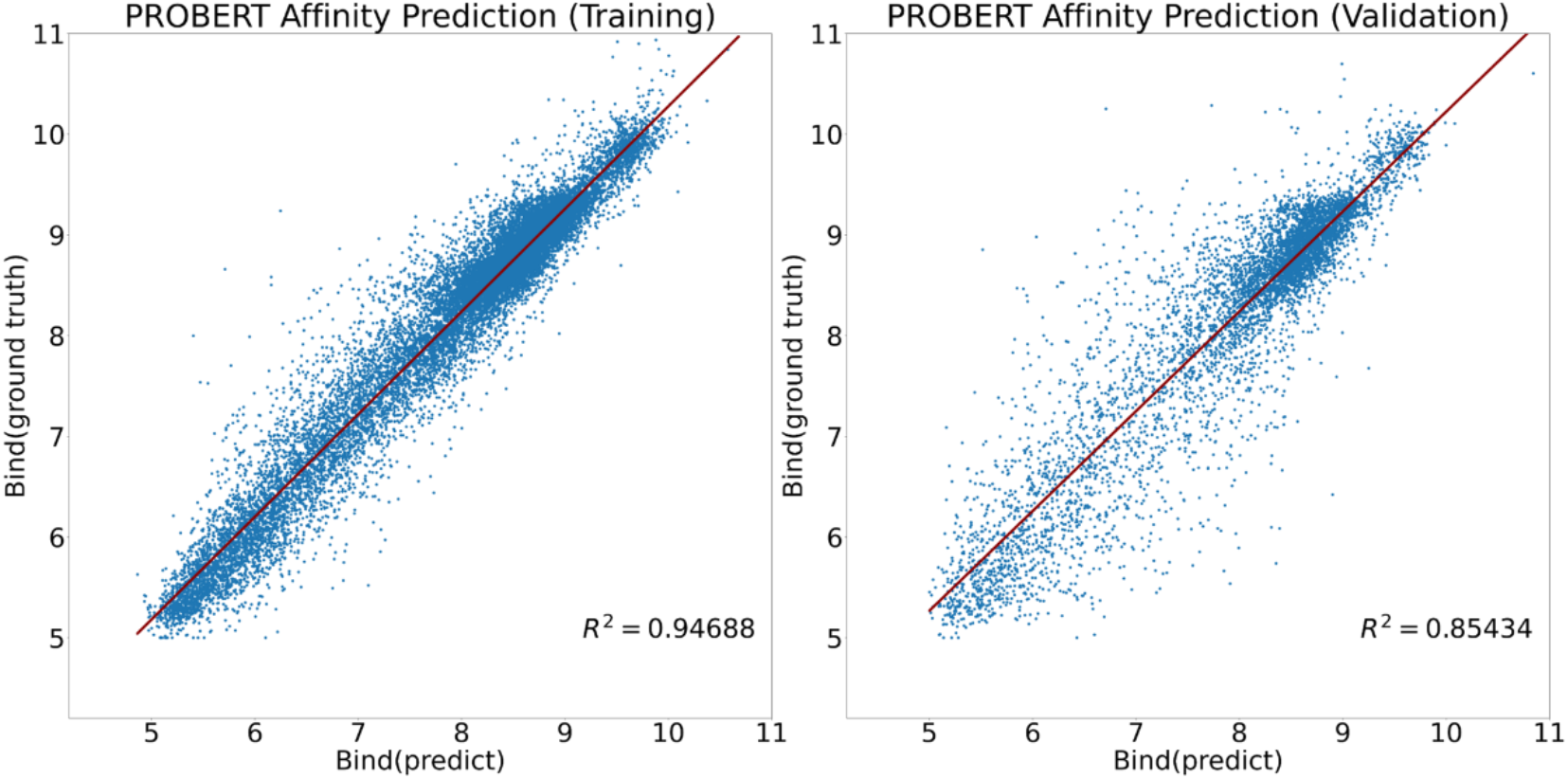
Affinity prediction results of ProBERT-VirEvol model

**Figure 7.**
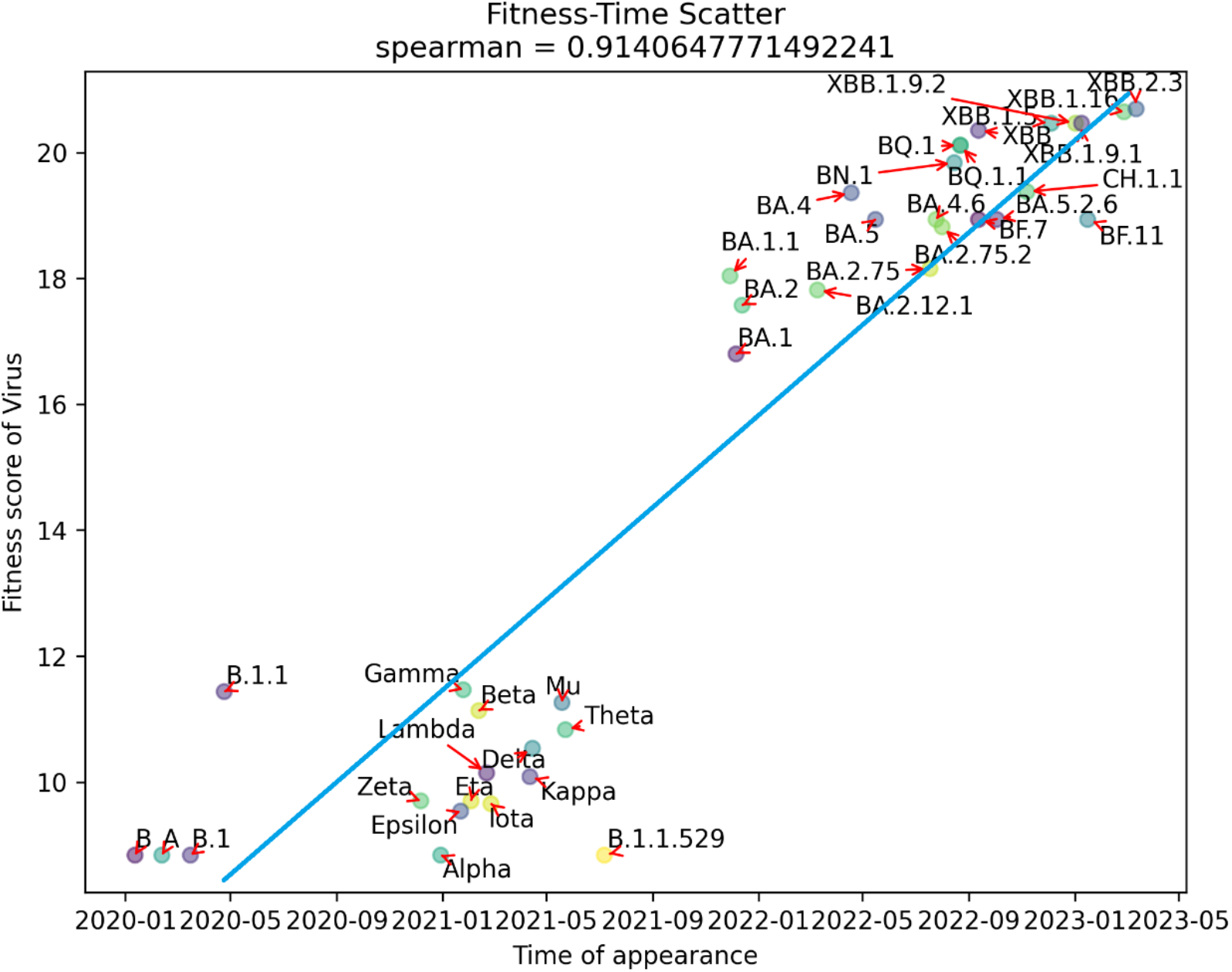
VirEvol model epidemic trend prediction, showing the correlation between the emergence time of COVID-19 strains and adaptation

**Figure 8.**
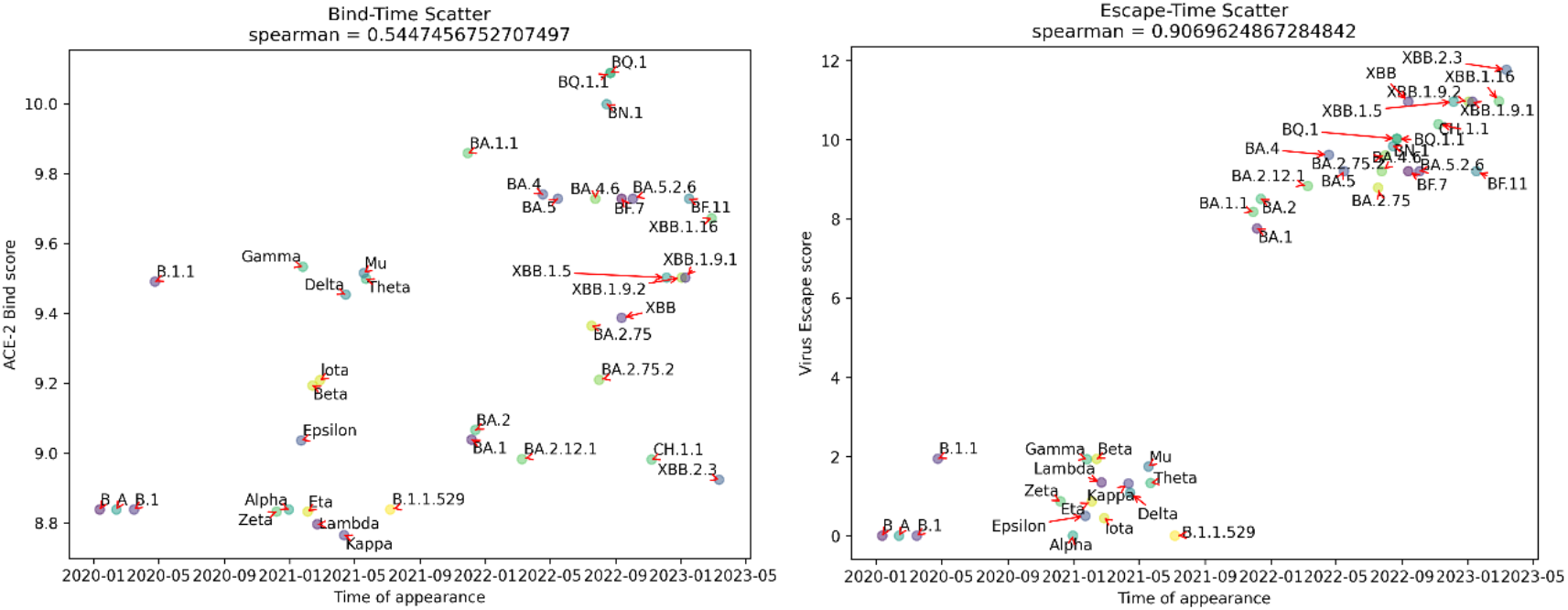
VirEvol model predicted the correlation between the emergence time of novel coronavirus strain and its affinity and immune escape score(the left figure is affinity and the right figure is immune escape score)

**Figure 9.**
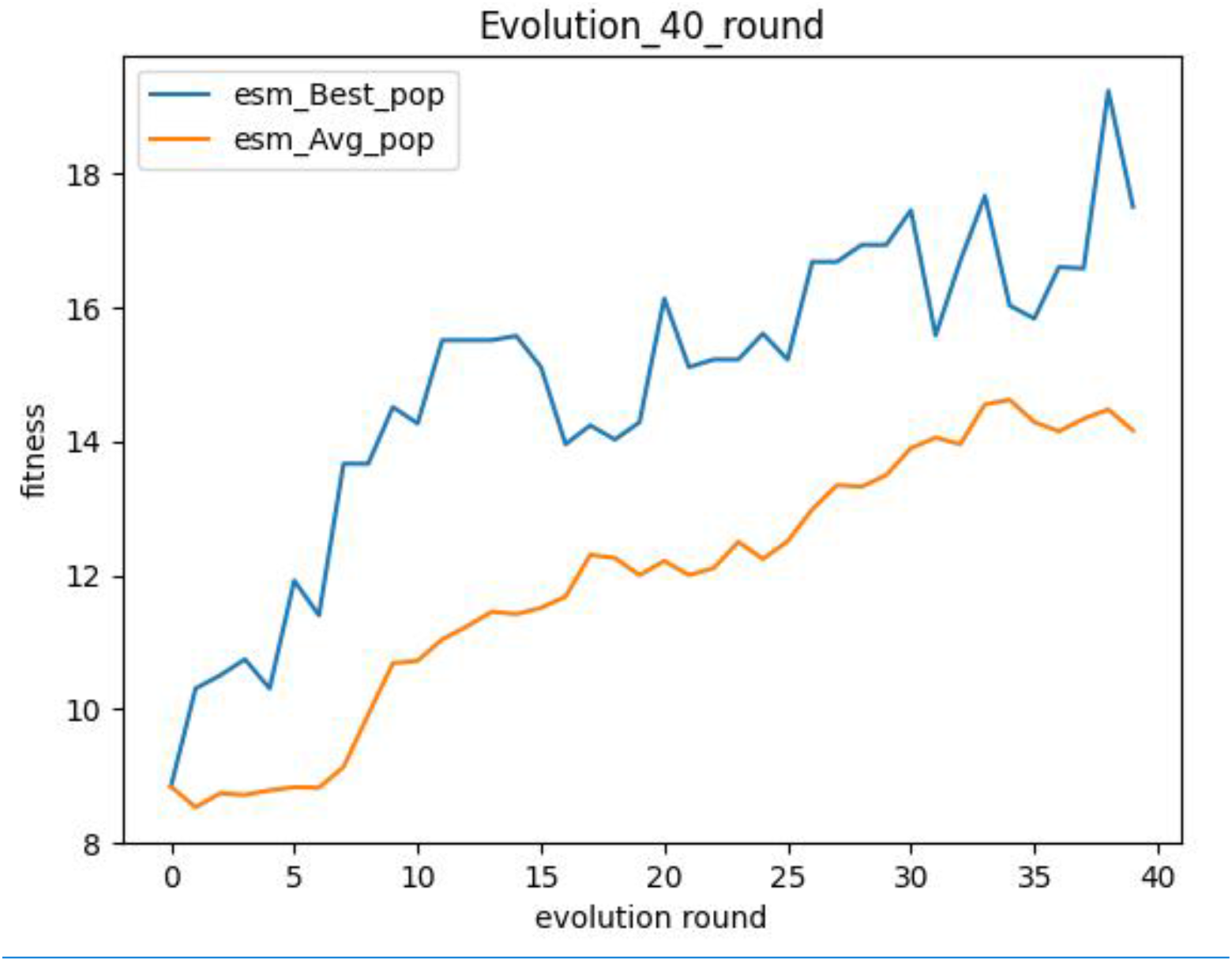
Exploring the genetic evolution of viruses (Blue represents the maximum fitness in the population, and yellow represents the average fitness in the population)

The two components of the fitness score, the correlation of affinity and immune escape score with the evolutionary direction of the virus, were calculated separately. The results showed that the correlation between virus evolution and target affinity was weak, and the Spearman correlation was only 0.54, while the Spearman correlation between immune escape and virus evolution was as high as 0.9. Therefore, the main evolutionary direction of virus evolution is immune escape, while maintaining a high affinity for ACE2 receptor (affinity score maintained above 8.8) during evolution.

### VirEvol uses genetic algorithms to explore the evolution of viruses

The built-in genetic algorithm can search for virus strains with higher adaptability based on the initial virus, with certain probabilities of gene recombination and gene mutation, exploring the future directions of virus mutations. As the number of iterations increases, the overall adaptability of the virus sequence continues to improve, as shown in Figure 9.

### VirEvol Predictions Reveal Molecular mechanisms of viral Evolution

Using VirEvol to predict and analyze the affinity and immune evasion scores of the XBB virus series, as shown in Figure 10, it is evident that XBB strains possess unique immune evasion scores higher than those of other lineages, consistent with reported results. This reveals one of the significant reasons for their ability to escape existing immune defenses and rapidly spread.

**Figure 10.**
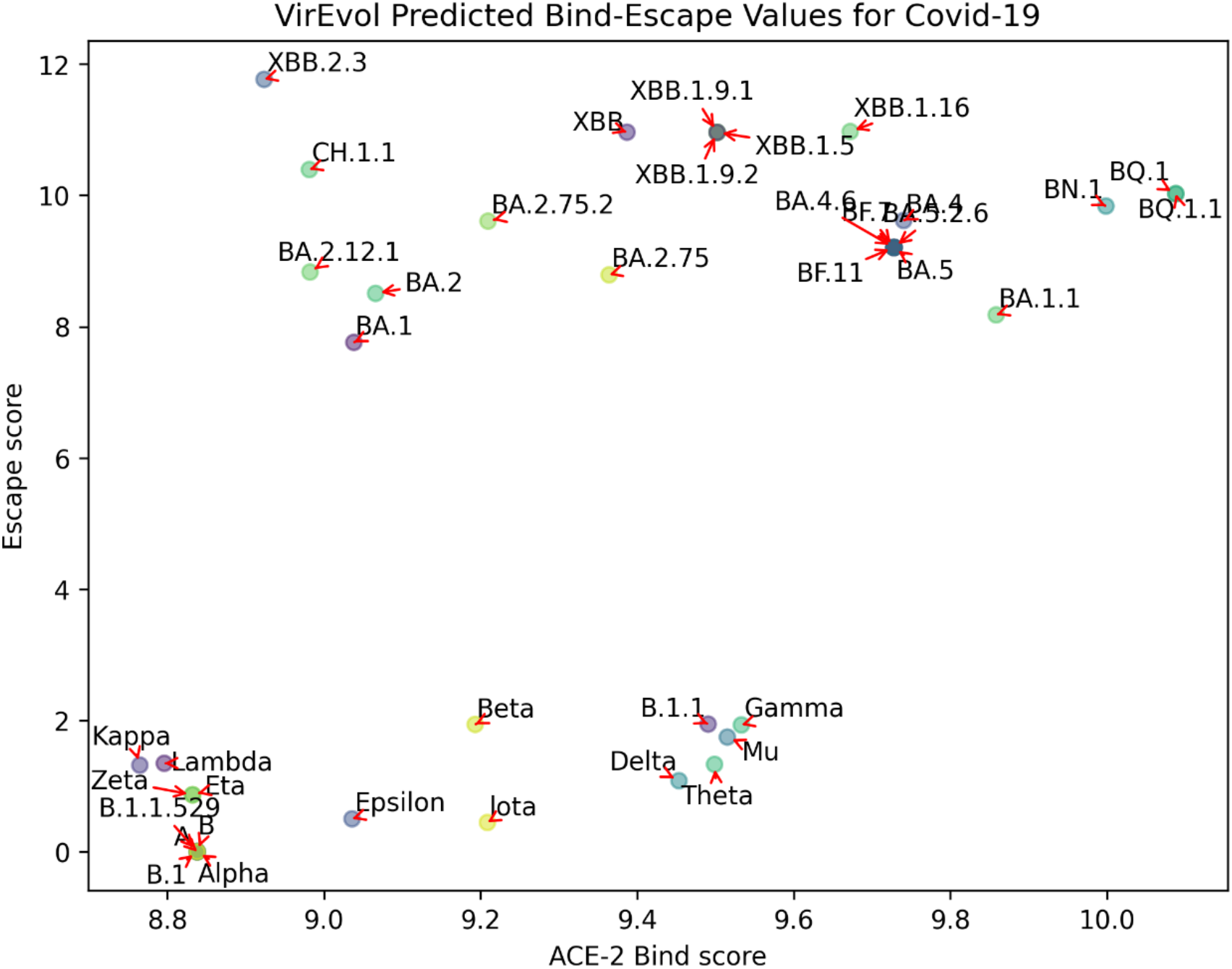
Affinity and immune escape scores of COVID-19 strains predicted by VirEvol model

By extracting the immune evasion scores of the XBB virus series from the model and combining them with multiple sequence alignments, we identified the key residues and their mutation mechanisms in XBB strains. Among these, the most significant contributors to immune evasion are the four sites: E484A, F486P, F490S, and Y505H. Research by Professor Andrew Pekosz from the Public Health of Johns Hopkins University School found that in subsequent strains like XBB.1.5, the F486P mutation replaced the original F486S in the original XBB. This mutation not only conferred immune evasion characteristics to XBB.1.5 but also enhanced the virus’s binding capacity to ACE2, thereby increasing its transmission ability.

## Conclusion

In this work, we proposed VirEvol, an evolutionary prediction platform for SARS-CoV-2 based on protein language modeling and structure and immune recognition mechanism, which can predict SARS-CoV-2’s affinity with the target ACE2 receptor and its immune escape ability, as well as the viral mutation trend. We collected a large amount of sequence data, deep mutation scanning data and virus-antibody complex structure data, and used a protein language model with 650 million parameters for feature extraction and task training. Our model achieved excellent performance on both affinity prediction and immune escape prediction subtasks. We also designed an adaptive score that integrates the information of both affinity and immune escape, thus revealing the molecular evolutionary mechanism of SARS-CoV-2. We found that SARS-CoV-2 mainly evolved with the goal of improving immune escape ability, while maintaining a high affinity level at the same time. This paper also analyzed the immune evasion mechanism of the XBB strains, and found that four sites, including F486P, contributed the most to it, and F486P also enhanced the binding ability of the virus to the ACE2 receptor and its transmission ability. In addition, this paper constructed an early warning system and a genetic evolution system, which accomplished early warning for 13 out of 14 high-risk strains with a success rate of 93%, and the genetic evolution system can carry out evolutionary exploration based on the initial strain RBD sequences.

The testable model of this paper provides artificial intelligence and data support for the construction of epidemic prevention and control system, and also provides important guidance for the over-design of vaccines and drugs for new SARS-CoV-2 mutants. Meanwhile, the overall architecture of the model, based on the design of protein language model and structure and immune recognition mechanism, as well as the prediction indexes combining affinity and immune escape also provide research ideas and technical support as reference for the prediction of evolution of other highly mutable viruses.

## Funding& Acknowledgements

B. Ma thanks support from Natural Science Foundation of China (Grant No. <u>32171246</u>) and Shanghai Municipal Government Science Innovation grant <u>21JC1403700</u>. Y. Wang thanks support from the grants from the National Natural Science Foundation of China (No.<u>32200531</u>), the Joint Research Funds for Medical and Engineering and Scientific Research at Shanghai Jiao Tong University (<u>YG2022QN114</u> and <u>YG2022QN082</u>), and Startup Fund for Young Faculty at SJTU (SFYF at SJTU). All computation was performed using high performance computer cluster of Shanghai Jiao Tong University.

## Reference

1. Jangra, S., et al., SARS-CoV-2 spike E484K mutation reduces antibody neutralisation. Lancet Microbe, 2021. 2(7): p. e283–e284.

2. Chen, Y., et al., Emerging SARS-CoV-2 variants: Why, how, and what’s next? Cell Insight, 2022. 1(3): p. 100029.

3. Hoffmann, M., et al., SARS-CoV-2 Cell Entry Depends on ACE2 and TMPRSS2 and Is Blocked by a Clinically Proven Protease Inhibitor. Cell, 2020. 181(2): p. 271–+.

4. Greaney, A.J., et al., Mapping mutations to the SARS-CoV-2 RBD that escape binding by different classes of antibodies. Nature Communications, 2021. 12(1).

5. Frank, F., et al., Deep mutational scanning identifies SARS-CoV-2 Nucleocapsid escape mutations of currently available rapid antigen tests. Cell, 2022. 185(19): p. 3603–+.

6. Starr, T.N., et al., Deep Mutational Scanning of SARS-CoV-2 Receptor Binding Domain Reveals Constraints on Folding and ACE2 Binding. Cell, 2020. 182(5): p. 1295–+.

7. Rives, A., et al., Biological structure and function emerge from scaling unsupervised learning to 250 million protein sequences. Proceedings of the National Academy of Sciences of the United States of America, 2021. 118(15).

8. Hsu, C., et al., Learning protein fitness models from evolutionary and assay-labeled data. Nature Biotechnology, 2022. 40(7): p. 1114–+.

9. Chen, C., et al., Computational prediction of the effect of amino acid changes on the binding affinity between SARS-CoV-2 spike RBD and human ACE2. Proceedings of the National Academy of Sciences of the United States of America, 2021. 118(42).

10. Wang, E. and A.K. Chakraborty, Design of immunogens for eliciting antibody responses that may protect against SARS-CoV-2 variants. Plos Computational Biology, 2022. 18(9).

11. Abbasi, W.A., et al., ISLAND: in-silico proteins binding affinity prediction using sequence information. Biodata Mining, 2020. 13(1).

12. Hie, B., et al., Learning the language of viral evolution and escape. Science, 2021. 371(6526): p. 284–+.

13. Wang, E., Prediction of antibody binding to SARS-CoV-2 RBDs. Bioinform Adv, 2023. 3(1): p. vbac103.

14. Ayijiang, Y., et al., Repeated Omicron infection alleviates SARS-CoV-2 immune imprinting. bioRxiv, 2023: p. 2023.05.01.538516.

15. Elnaggar, A., et al., ProtTrans: Toward Understanding the Language of Life Through Self-Supervised Learning. IEEE Trans Pattern Anal Mach Intell, 2022. 44(10): p. 7112–7127.

16. Meier, J., et al., Language models enable zero-shot prediction of the effects of mutations on protein function. Advances in Neural Information Processing Systems 34 (Neurips 2021), 2021. 34.

17. Beguir, K., et al., Early computational detection of potential high-risk SARS-CoV-2 variants. Computers in Biology and Medicine, 2023. 155.

18. Taft, J.M., et al., Predictive profiling of SARS-CoV-2 variants by deep mutational learning. bioRxiv, 2021: p. 2021.12.07.471580.

19. Gowthaman R, Guest JD, Yin R, Adolf-Bryfogle J, Schief WR, Pierce BG. CoV3D: a database of high resolution coronavirus protein structures. Nucleic Acids Res. 2021;49(D1):D282–D287.

20. Wang M, Zhu D, Zhu J, Nussinov R, Ma B. Local and global anatomy of antibody-protein antigen recognition. J Mol Recognit. 2018;31(5):e2693.

